# Prenatal cannabinoid exposure induces sex-specific alterations in placental growth and lipid metabolism gene expression

**DOI:** 10.64898/2026.06.24.734289

**Authors:** Rachel C. West, Adrian H. Courville, Cristine R. Camp, Paige Drotos, Carly Parker, Miranda N. Reed

**Affiliations:** Anatomy, Physiology, Pharmacology Department, College of Veterinary Medicine, Auburn University; Department of Drug Discovery and Development, Harrison College of Pharmacy, Auburn University

**Keywords:** Cannabis, pregnancy, metabolism, placenta

## Abstract

**Background:** Prenatal cannabis use is becoming increasingly more commonplace. However, cannabis exposure is linked to adverse pregnancy outcomes, including gestational hypertension, preeclampsia, and preterm birth. The aim of this study was to determine the morphological and molecular effects of prenatal cannabinoid exposure on the placenta.

**Methods:** Pregnant Sprague-Dawley rats were exposed daily to vaporized THC (100 mg/mL) starting at gestational day (GD)5 until GD19 when dams were sacrificed and fetuses and placentas collected. Fetuses were genotyped for genetic sex and transcriptomic analysis was performed on male and female THC-exposed and control placentas.

**Results:** On GD19, both the fetuses and placentas from the THC group were significantly larger than the control. When separated by sex, both male and female THC fetuses were significantly larger; however, only male THC placentas were significantly larger than male control placentas with no significant difference in placental weight between female control and THC placentas. RNA-sequencing revealed enriched biological processes related to nutrient transport and lipid catabolism, protein-lipid complex formation, and lipoprotein particle remodeling and organization. Further transcriptomic analysis determined that the differentially expressed genes and enriched biological processes related to lipid metabolism were preferentially enriched in the female THC placentas compared to the male, suggesting a sex-specific effect.

**Discussion:** Collectively, these data present sex-specific effects of prenatal cannabinoid exposure on placental growth and global gene expression. These data also suggest that sex influences gene expression of genes related to lipid metabolism in the THC-exposed placentas.

## Introduction

Cannabis use during pregnancy has increased substantially in the United States. Recent data from the Centers for Disease Control and Prevention indicate that 6.6% of pregnant women reported cannabis use within the past 30 days and 4.2% reporting near daily use^1^. However, toxicological screening for cannabis in other studies suggests that prenatal cannabis exposure may be considerably underreported, with metabolites of cannabis detected in 15-23% of pregnant women^2, 3^. Because there are limited options for symptom relief during pregnancy, women often use cannabis to self-treat nausea, insomnia, chronic pain, and anxiety^4, 5^. This practice may also be driven by the perception that cannabis use carries little to no risk^4^. However, prenatal cannabis use and gestational hypertension, preeclampsia, and preterm birth are linked^6, 7^.

Given these risks, it is critical to understand how cannabis affects maternal-fetal physiology, particularly at the level of the placenta. During pregnancy, the placenta serves as a protective barrier between maternal insults and fetal health and development. Despite this role, the placenta is particularly vulnerable to tetrahydrocannabinol (THC), the primary psychoactive component of cannabis. THC can readily cross the placenta from the maternal circulation and enter the fetal blood stream^8^. Additionally, the placenta is enriched with cannabinoid receptors; CB1 and CB2 are present in all placental cell subtypes as early as the first trimester^9^.

Experimental studies using rat models have demonstrated that prenatal THC exposure can induce adverse changes in placental morphology^10, 11^. Notably, THC-exposed placentas exhibit reduced levels of glucose transporter 1 (GLUT1) and glucocorticoid receptor (GR)^11^, suggesting prenatal cannabinoid exposure can potentially alter metabolism and nutrient transport between the mother and fetus.

The relationship between placental and fetal metabolism is intimately interconnected. The placenta must consume maternal nutrients to provide energy for its function but the placenta also serves as a conduit, transporting essential nutrients to the fetus^12^. There is also convincing evidence that prenatal cannabinoid exposure influences fetal metabolism. Children born from mothers who used cannabis while pregnant have a higher body mass index (BMI), increased adiposity, elevated fasting glucose levels, and higher triglyceride levels^13–15^.

In this study, we used a rat model of prenatal cannabinoid exposure to assess changes in placental and fetal size and placental global gene expression. We exposed pregnant rats daily to a dose of vaporized THC beginning shortly after embryo implantation and lasting until late gestation. We used bulk RNA-sequencing (RNAseq) to assess differences in gene expression and found evidence that supports the hypothesis that prenatal exposure to THC can alter placental metabolism. We also assessed the differences in male and female placentas and found that fetal sex influences how the placenta responds to THC exposure.

## Materials and Methods

### Animals

Timed pregnant Sprague-Dawley rats were purchased from Envigo Laboratories and arrived to Auburn University on gestational day (GD)3. Pregnant rats (n = 3) were allowed to acclimatize to their new environment for 48 hours and began receiving a daily dose of delta-9-tetrahydrocannabinol (THC, 100 mg/mL) vaporized in polyethylene glycol 400 (PEG) from GD5 to GD19. Vehicle control (VEH) pregnant rats (n = 4) were exposed to daily vaporized doses of PEG from GD5 to GD19. Pregnant rats were euthanized on GD19, and fetectomies performed.

Animals were housed in a controlled vivarium with regulated temperature and humidity maintained in a 12 hour light/dark cycle. Animals were provided food and water ad libitum. All care and procedures were performed within compliance of the National Institutes of Health (NIH) Care and Use of Laboratory Animals guidelines and approved by the Auburn University Animal Care and Use Committee (IACUC).

### Tissue collection and preparation

Pregnant dams were euthanized using carbon dioxide followed by decapitation at GD19. The uterus was removed, then each fetus and placenta was dissected and weighed. Placentas and fetal tail snips were collected then snap-frozen in liquid nitrogen for further assessment.

Fetal sex was determined by genotyping using the fetal tail snips as previously published^16^. Briefly, DNA was isolated from tail snips using DNA lysis buffer (0.3% Nonidet P-40 substitute, 50 mM Potassium chloride, 10 mM Tris, 0.3% Tween20, and 1 mg/mL Proteinase K). Tail snips were incubated in lysis buffer at 55°C for 1 hour followed by 98°C for 15 minutes. After incubation, DNA was centrifuged at 2,000 x g for 3 minutes then supernatant was collected and DNA quantified. DNA was diluted to 50 ng/µL then added to 2X Phusion U Green Supermix (FisherScientific, #F564S). Primers for the X- and Y-chromosome homologs *DDX3X* and *DDX3Y* were added to PCR supermix and multiplex PCR performed using the following PCR protocol of 98°C for 30 seconds, then 40 cycles of 98°C for 10 seconds, 58°C for 30 seconds, and 72°C for 42 seconds. The PCR was completed with a final incubation of 72°C for 5 minutes then PCR product collected for gel electrophoresis. After electrophoresis, PCR gels were visualized and samples that had bands for *DDX3X* and *DDX3Y* were considered male and samples that only had bands for *DDX3X* considered female.

### RNA isolation and RNA-sequencing

Four to six placentas from each dam were chosen at random from the THC (n = 20, 10 male, 10 female) and VEH (n = 20; 10 male, 10 female) group were chosen at random for RNA isolation and bulk RNA-sequencing (RNAseq). Total RNA was isolated from placentas using the Qiagen RNeasy mini kit (#74104) following manufacturer’s instructions. Purified RNA was sent to Novogene for library construction and sequencing. Libraries were pooled and sequenced using the Illumina NovaSeq X Plus platform.

Sequencing yielded a mean 59,477,936 (standard deviation = 7,275,649) raw read count from the VEH group and 57,869,882 (standard deviation = 9,271,476) raw read count from the THC group. Raw fastq reads were subject to quality control using fastp and adapters and low-quality reads were removed. Cleaned reads were mapped to the Rattus norvegicus (mRatBN7.2) reference genome. Individual reads were also adjusted to provided fragments per kilobase per transcript sequence per millions base pairs sequenced (FPKM) values. The DESeq2 R package (1.42.0) was used to perform differential expression analysis and genes that met the threshold of an adjusted p-value (padj) of < 0.05 and a log2foldchange >0.5 were considered statistically significant. Gene ontology (GO) and Kyoto Encyclopedia of Genes and Genomes (KEGG) enrichment analysis of differentially expressed genes was performed using the clusterProfiler R package (4.8.1). GO terms with a padj of <0.05 were considered significantly enriched. For additional analysis of DEGs, enrichment analyses, and visualization of data, we used the NovoMagic free Novogene platform.

### Statistics

We first assessed fetal and placental weight data for normality using the Shapiro-Wilk test, which indicated that both the control and treatment groups deviated significantly from a normal distribution. The control group yielded a Shapiro-Wilk statistic of W = 0.9394 (p = 0.0141), while the treatment group yielded W = 0.3802 (p < 0.0001). Since the assumption of normality was not met, we performed nonparametric statistical analyses, specifically Mann-Whitney U tests to assess differences between groups. P-values < 0.05 were considered statistically significant. Statistical analyses were conducted using the GraphPad Prism for Windows software (Version 11.0.0).

## Results

### Vaporized THC exposure results in increased placental and fetal weight in a sex-specific manner

We first evaluated litter size, fetal weight, placental weight, and the fetal:placental weight ratio. THC did not affect litter size; however, THC fetuses and placentas were both significantly (p < 0.01) larger than the VEH group (Table 1, Figure 1A-B). We next assessed if the differences in fetal and placental weights were influenced by sex. Both male and female THC fetuses were significantly larger than their VEH counterparts (p < 0.01, Figure 1D); however, only the male placentas were significantly larger than the male VEH placentas (p < 0.001, Figure 1C) with no significant difference in female placental weights (Table 1). Further analysis assessing the fetal:placental weight ratio revealed that THC exposure had no effect on fetal:placental weight ratio (Table 1).

**Figure 1.**
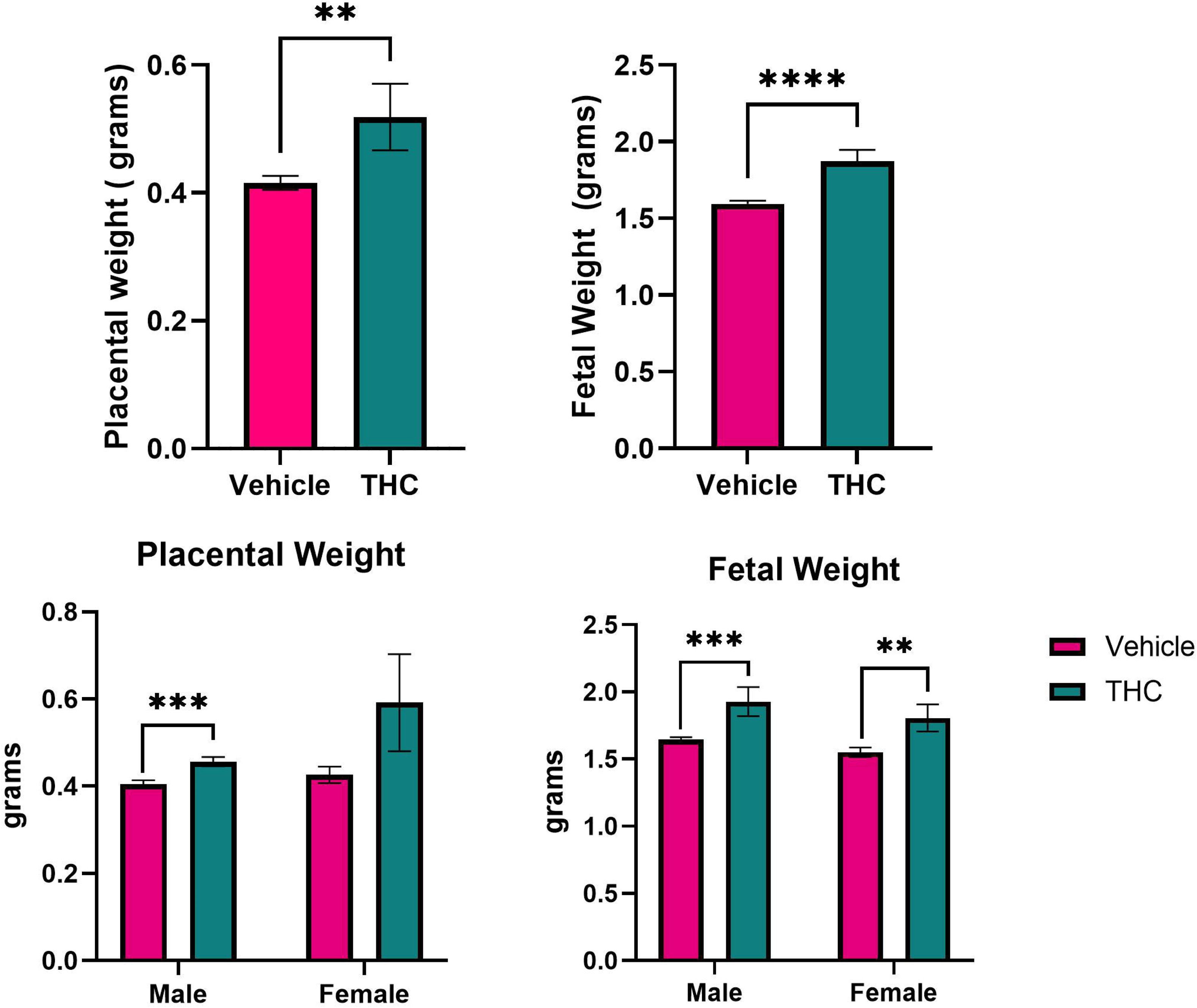
Prenatal cannabinoid exposure increases placental and fetal weights. Placental (A) and fetal (B) weights were significantly increased in GD19 THC-exposed pregnancies. When comparing weights between the sexes, only male THC placentas were significantly heavier than male controls with no significant difference between female THC and female control placentas. Both male and female fetuses were significantly larger than their control counterparts. Values are mean ± SEM. n = 49 placentas (from 4 dams) for the VEH group and n = 39 placentas (from 3 dams) from the THC group. For statistical comparison between sexes n = 23 VEH males and 26 VEH females and 21 THC males and 18 THC females. Statistics were generated using the Mann-Whitney U test. ** p < 0.01, *** p < 0.001 **** p < 0.0001.

**Table 1.** Fetal and placental outcome measurements. Litter size, fetal weight, placental weight, and fetal:placental weight ratios in VEH, THC, and male and female VEH and THC groups. n = 49 VEH (4 dams) and n = 39 THC (3 dams). For sexes, n = 23 male VEH, n = 26 female VEH, n = 21 male THC and n = 18 female THC. Values are mean ± SEM. P-values generated using Mann-Whitney U test.

|  | VEH | THC | p-value |
| --- | --- | --- | --- |
| Litter size | 12.25±1.109 | 13±0.5774 | p = 0.578 |
| Fetal weight | 1.595±0.021 | 1.872±0.074 | p < 0.0001 |
| Placental weight | 0.415±0.011 | 0.519±0.052 | p = 0.003 |
| Fetal:placental weight ratio | 3.96±0.111 | 4.04±0.190 | p = 0.931 |
| Male fetal weight | 1.65±0.0189 | 1.93±0.108 | p = 0.007 |
| Male fetal:placental weight ratio | 4.12±0.106 | 4.29±0.281 | p = 0.499 |
| Male placental weight | 0.404±0.009 | 0.456±0.011 | p = 0.0009 |
| Female fetal weight | 1.55±0.034 | 1.81±0.102 | p = 0.003 |
| Female placental weight | 0.426±0.019 | 0.592±0.112 | p = 0.283 |
| Female fetal:placental weight ratio | 3.82±0.185 | 3.75±0.239 | p = 0.75 |

### Vaporized THC exposure induces sex-specific global transcriptional changes

We next used RNAseq to identify differentially expressed genes (DEGs) in the placentas. After setting the threshold of significance at padj < 0.05 and a log2foldchange > 0.5, we identified 480 DEGs (358 up and 122 down) in the THC group compared to the VEH (Figure 2A, Supplemental File 1). Interestingly, when we compared DEG expression between the sexes we found differing patterns between the male and female groups. There were only 44 DEGs (33 up and 11 down) in the THC males versus VEH males (Figure 2B, Supplemental File 1). Comparatively, there were 259 DEGs (177 up and 82 down) in the THC female placentas compared to the VEH female placentas (Figure 2C, Supplemental File 1). We next compared the DEG profiles between the male THC vs. VEH placentas and the female THC vs. VEH placentas to identify genes and pathways that were preferentially enriched by sex. This analysis identified 140 DEGs preferentially enriched in the female THC placentas and 21 DEGs preferentially enriched in the male THC placentas (Figure 2D, Supplemental File 2). Hierarchical clustering analysis was then performed to visualize differences in expression patterns among these preferentially enriched genes (Figure 2E). These results indicate that THC exposure alters placental gene expression in both sexes; however, the effects are more extensive and pronounced in female placentas.

**Figure 2.**
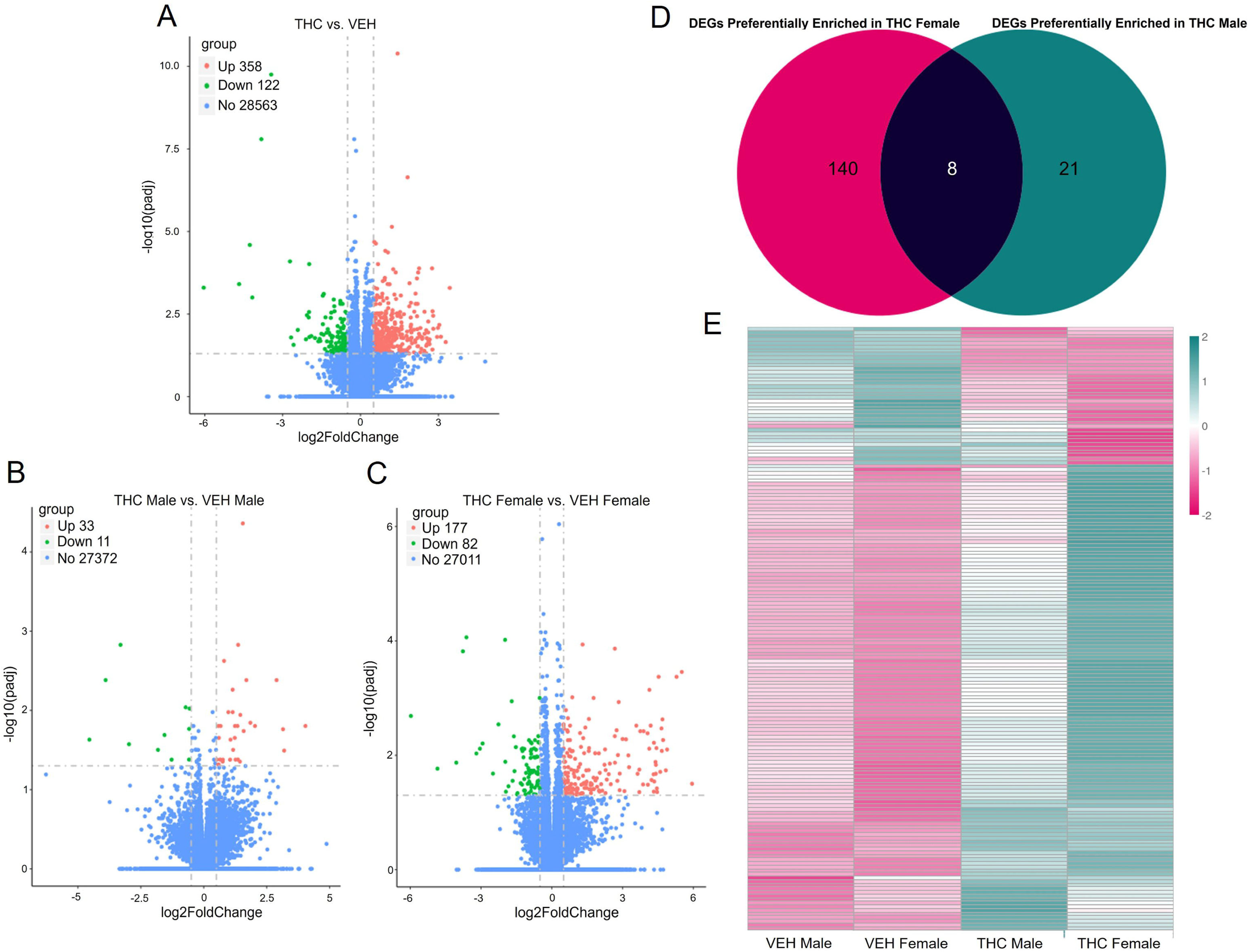
Prenatal cannabinoid exposure induces global gene expression differences in the placenta. A) There were 480 (358 up, 122 down) differentially expressed genes between the THC and VEH placentas (n= 20). When comparing between the sexes (n=10), B) THC males had a smaller range of DEGs with only 44 DEGs (33 up, 11 down) compared to VEH males whereas C) THC females had 259 DEGs (177 up, 82 down) compared to VEH female placentas. D) Comparison between the two DEG datasets indicated that there were 140 DEGs preferentially enriched in the THC females compared to THC males, VEH females, and VEH males. Alternatively, there were 21 DEGs preferentially enriched in the THC male placentas compared to THC females, VEH females, and VEH males. E) Clustering analysis was performed to visualize the differences in gene expression of the male and female preferentially enriched genes. Significance was determined if genes were below the threshold of an adjusted p-value < 0.05 and above the threshold of a log2foldchange > 0.5.

### GO enrichment analysis reveals THC exposure alters pathways related to lipid metabolism

GO enrichment analysis of DEGs between the THC and VEH groups identified several significantly enriched pathways related to lipid catabolism, regulation of lipoprotein levels, and protein-lipid complex remodeling (Figure 3A, Table 2, Supplemental File 3). In addition, pathways associated with wound healing, monocyte and lymphocyte chemotaxis, superoxide metabolism, and acute inflammatory response were significantly enriched in the THC-exposed placentas (Supplemental File 3), suggesting increased inflammation and cellular stress.

**Figure 3.**
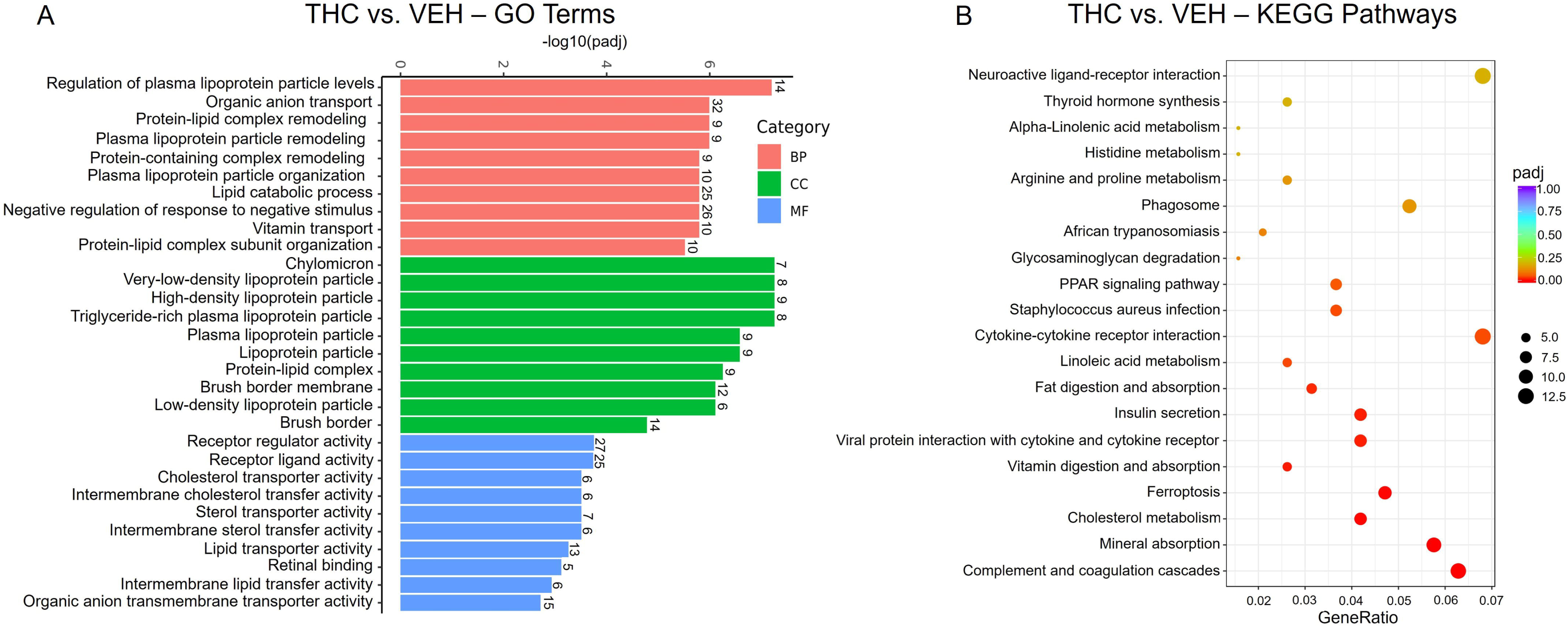
Pathway enrichment analysis of DEGs in THC placentas reveal alterations in pathways related to lipid metabolism, nutrient transport, and cellular stress. A) Gene ontology of THC DEGs identified several GO terms related to lipid catabolism, lipoprotein organization, nutrient transport, and cytokine binding. B) KEGG analysis identified KEGG terms related to metabolism, absorption, insulin secretion, and cellular stress were significantly enriched in the THC placentas. GeneRatio represents the proportion of DEGs mapped to a specific pathway over the total size of the input gene list. Dot size refers to the absolute count of enriched genes. Terms were considered significantly enriched if they had an adjusted p-value < 0.05.

**Table 2.**
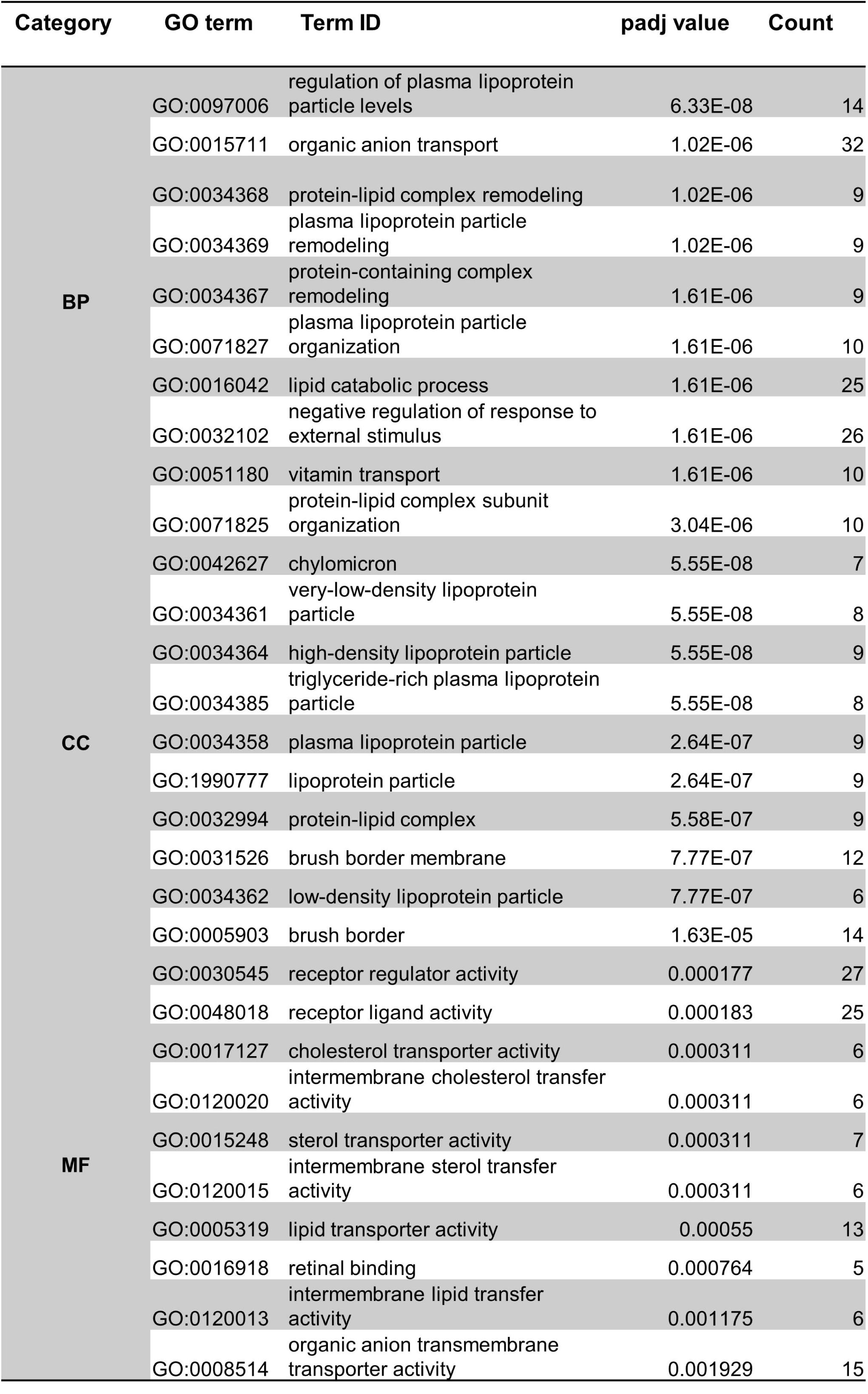
Top 10 GO terms enriched in THC placentas. The top 10 GO terms separated by category; Biological Process (BP), Molecular Function (MF), and Cellular Component (CC). Count refers to the number of DEGs that are annotated to each specific GO term. A comprehensive list of all significantly enriched GO terms can be found in Supplemental File 3.

To further characterize the biological functions of DEGs in the THC-exposed placentas, we performed KEGG pathway enrichment analysis. Many of the top enriched KEGG pathways were related to metabolism and nutrient absorption (Figure 3B, Supplemental File 3).

Specifically, pathways involved in cholesterol metabolism, insulin secretion, fat digestion and absorption, mineral absorption, vitamin digestion and absorption, and linoleic acid metabolism were significantly enriched in THC placentas. Pathways associated with cellular stress and immune signaling including ferroptosis, viral protein interaction with cytokine and cytokine receptor, and cytokine-cytokine-receptor interaction were also significantly enriched (Figure 3B).

We also performed GO on the DEGs preferentially enriched in the female THC placentas. Interestingly, the majority of the significantly enriched GO terms enriched were also related to lipid catabolism, lipoprotein remodeling and organization, and protein-lipid complex assembly (Figure 4A, Table 3, Supplemental File 3). To visualize expression of patterns of genes within these enriched pathways, we conducted hierarchical clustering analysis across male and female VEH and THC groups (Figure 4B). Many of the genes contributing to these pathways belonged to the apolipoprotein (Apo) family, including *Apob, Apoa2, Apoh, Apoc2, Apoa1, Apom,* and *Apoe.* Additional genes represented within the enriched GO terms included retinol binding protein 4 (*Rbp4*), solute carrier family 27 member 2 (*Slc27a2*), phospholipase A2 group XIIB (*Pla2g12b*), ghrelin and obestatin prepropeptide (*Ghrl*), and retinol binding protein receptor 2 (*Rgd1305807*).

**Figure 4.**
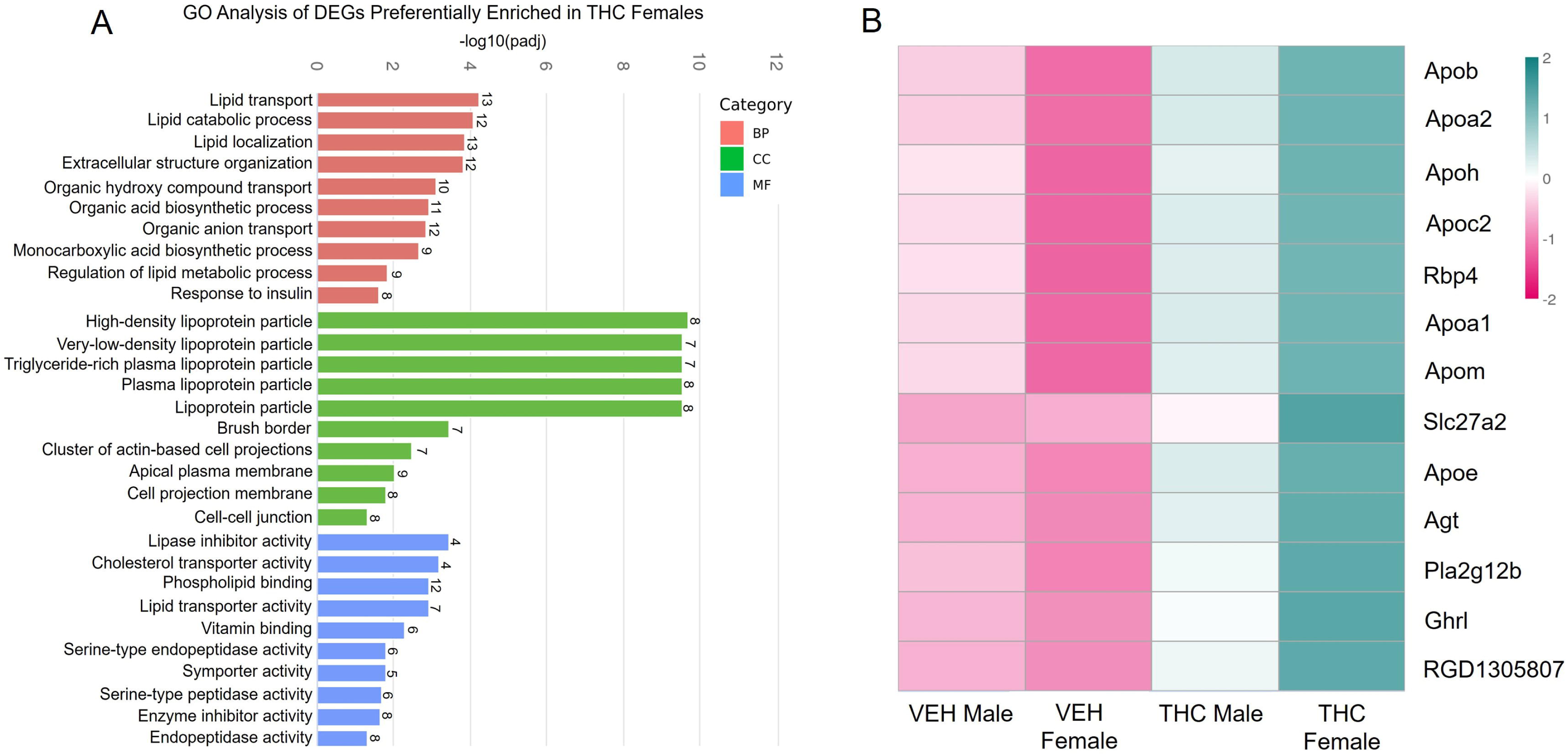
GO analysis of DEGs preferentially enriched in THC females suggest that lipid metabolism is skewed in a sex-specific manner. A) GO performed on the DEGs preferentially enriched in the female THC placentas identified pathways related to lipid catabolism, transport, metabolism, and localization, suggesting that the alterations in lipid metabolism are augmented in female placentas compared to male. B) Clustering analysis between the 4 treatment groups of the top genes found in the lipid related GO terms. Many of the genes are a part of the Apo family. Terms were considered significantly enriched if they had an adjusted p-value < 0.05.

**Table 3.**
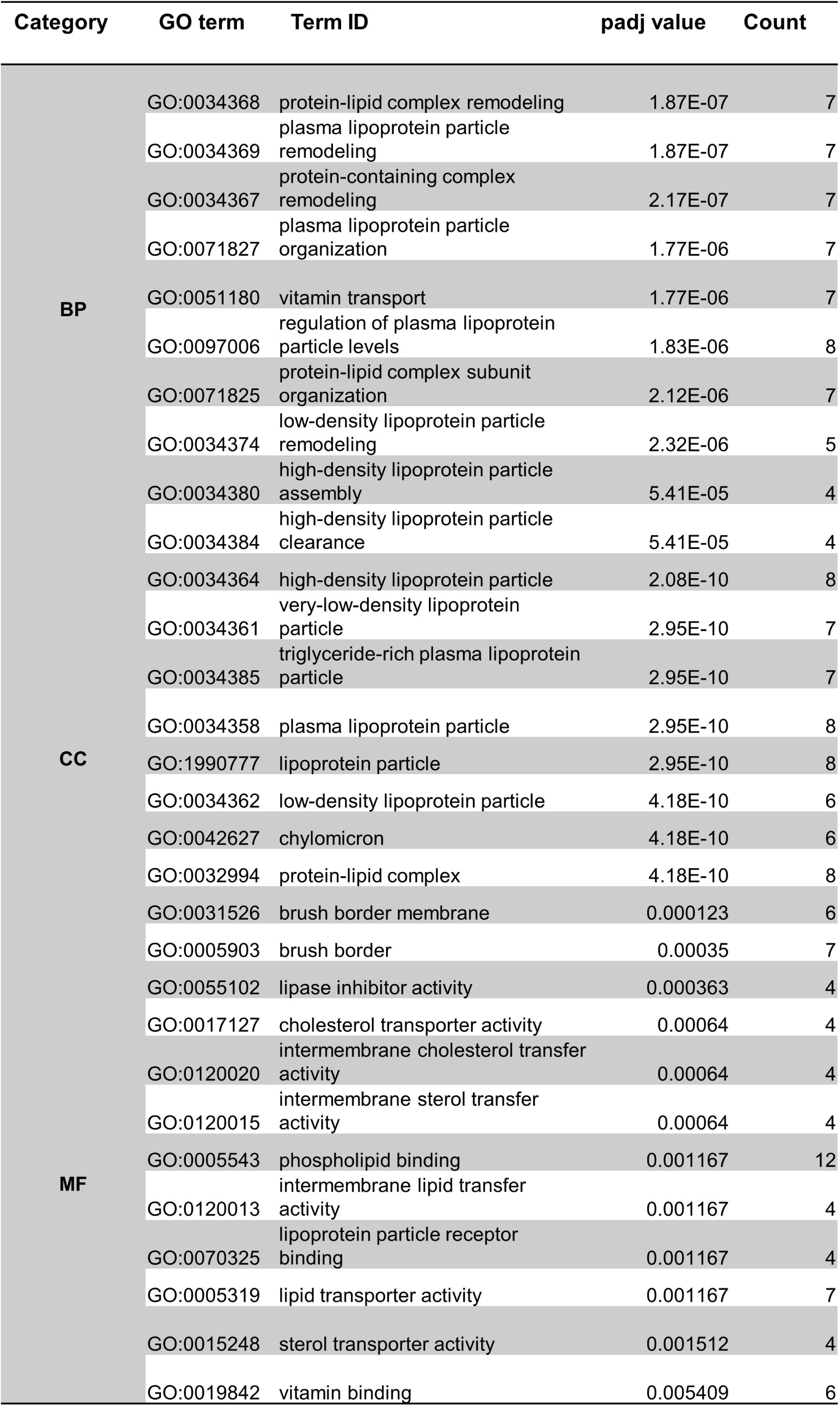
Top 10 GO terms identified from the preferentially enriched DEGs in female THC placentas. The top 10 GO terms separated by category; Biological Process (BP), Molecular Function (MF), and Cellular Component (CC). Count refers to the number of DEGs that are annotated to each specific GO term. A comprehensive list of all significantly enriched GO terms can be found in Supplemental File 3.

## Discussion

As the use of cannabis during pregnancy continues to rise, better understanding the physiological consequences on fetal and placental programming is critical. In this study, we provide global gene expression data demonstrating that prenatal cannabinoid exposure alters the transcriptome of the placenta in a sex-specific manner. Furthermore, many of the differentially expressed genes are related to lipid catabolism, lipid-protein complex formation, and lipoprotein particle remodeling and organization, suggesting that THC has the potential to alter placental lipid metabolism.

Using our rat model, we found that inhalation of vaporized THC during pregnancy increased both placental and fetal weights by GD19. This finding was unexpected as previous rat studies of prenatal cannabinoid exposure have reported reduced birth weight and lower offspring weight at postnatal day (PND)5^11, 17^. However, two studies that also assessed fetal weight between GD18-19.5 similarly observed modest increases in fetal weight following THC exposure, although these differences did not reach statistical significance^11, 18^. In contrast, both studies reported significantly reduced placental weights in the THC-exposed groups^11, 18^. Notably, the routes of THC administration differed between these studies, with one using intraperitoneal injection^11^ and the other exposing pregnant mice to cannabis smoke^18^.

Alternatively, another study using an edible THC dosing strategy reported a reduction in fetal weight by GD19.5^19^. These discrepancies suggest that the route of THC administration may contribute to variability in fetal and placental growth outcomes. It is also important to note that GD19 in the rat roughly corresponds to the late second to early third trimester of human pregnancy^20, 21^. Thus, the reported increase in placental and fetal weights at GD19 should also be interpreted within the broader context of the prenatal cannabinoid exposure studies that assess outcomes at birth and during postnatal development. Collectively, these findings support the hypothesis that prenatal cannabinoid exposure alters fetal and placental developmental programming.

We next used RNAseq, along with downstream enrichment and pathway analyses, to characterize the transcriptional signature of THC-exposed placentas compared to vehicle control placentas. Transcriptomic analysis revealed key differences in genes related to metabolism, nutrient transport, and insulin response in THC-exposed placentas. Additionally, these data suggest that prenatal cannabinoid exposure induces the expression of genes related to cellular stress, complimenting previous work performed in immortalized placental cell lines demonstrating that exposure to cannabinoids in vitro can induce cellular stress^22, 23^.

Cannabis use during pregnancy has been linked to preeclampsia, intrauterine growth restriction (IUGR), and preterm birth^24–26^; all complications that are often driven by placental insufficiency^27^. Previous studies have shown that pregnant women who use cannabis exhibit impaired placental angiogenesis, findings that are supported by mouse models of prenatal cannabinoid exposure^28^. Similarly, daily intraperitoneal injection of THC in pregnant rats has been shown to induce a phenotype representative of placental insufficiency and resulted in growth-restricted offspring^11^.

We propose that our transcriptome data provide potential evidence at the transcriptome level of placental metabolic dysfunction. The placenta acts as a nutrient sensor, actively responding to maternal and fetal metabolic signals^29^. It integrates multiple nutrient-sensing pathways to coordinate nutrient uptake and distribution to the fetus; however, these pathways can be impaired by cellular stress, leading to metabolic dysfunction and placental insufficiency^30^.

Consistent with this, several studies have reported altered lipid and energy metabolism in pregnancies complicated by IUGR^31–33^. In healthy pregnancies, high levels of lipids is a physiological adaptation that facilitates rapid synthesis of lipoproteins to meet the energy demands for fetal development^34^. However, in cases of placental insufficiency where uteroplacental blood flow is reduced, the placenta often exhibits excessive lipid catabolism and accumulation. This may represent a compensatory mechanism to sustain placental energy needs and an adequate supply of fatty acids to be delivered to the fetus^35^.

Many of the differentially expressed genes related to lipid metabolism belong to the apolipoprotein family. Apolipoproteins are structural and regulatory proteins that support lipid metabolism and transportation into both the maternal and fetal circulation during pregnancy^36^. The placenta actively synthesizes apolipoproteins and transports them to the maternal circulation to aid in cholesterol regulation during pregnancy^37^. Alternatively, placental apolipoproteins that are transported to the fetus aid in fetal development, growth, and neurodevelopment^38, 39^.

However, elevated levels of several apolipoproteins have been associated with preeclampsia^40–42^ and elevated apolipoprotein levels have been suggested as a potential biomarker for preeclampsia. While our transcriptomic data is descriptive in nature, it does provide evidence of a broader placental dysfunction that may be contributing to placental insufficiency.

Interestingly, when we analyzed the data while accounting for fetal sex, it appears that many of the DEGs related to lipid metabolism and transport and response to insulin are preferentially enriched in the THC female placentas compared to the THC male placentas. These data are consistent with previous reports that prenatal cannabinoid exposure induced female-specific glucose intolerance and insulin resistance in PND21 offspring^43^, suggesting that female placentas and offspring are more susceptible to the effects of prenatal cannabinoid exposure.

There are inherent sex-specific differences in fetal and placental metabolism with female placentas more responsive to their environments and having more flexible nutrient-sensing pathways^30, 44^. Our DEG data also supports this premise with considerably larger numbers of DEGs in the female THC placentas compared to female VEH placentas than the male THC placentas compared to male VEH placentas, suggesting again that female placentas are more responsive to their environment.

We recognize there are limitations of this study. First, we have a relatively small sample size for dams which may have limited the statistical power to detect modest associations. Consequently, some potentially meaningful relationships may not have reached statistical significance. Replication in larger cohorts will be helpful to confirm the robustness of these findings. Another limitation is the sole use of RNAseq to profile gene expression changes in the THC-exposed placentas. Although RNAseq provides a comprehensive and sensitive assessment of transcriptomic changes, biological validation through protein-level analyses would strengthen confidence in the observed gene expression patterns. Consequently, the results provided in this study should be interpreted as hypothesis-generating. Nevertheless, the observed trends were consistent the previous reports of placental dysfunction and provide helpful preliminary evidence that warrants further investigation in larger studies.

In summary, this study demonstrates that prenatal exposure to vaporized THC induces differences in placental gene expression in genes related to metabolism, nutrient transport, and insulin response. Furthermore, these differences are primarily driven by augmented expression of genes in female THC-exposed placentas. These differences in gene expression correlated with larger GD19 placental and fetal weights, suggesting a phenotypic difference in growth and development caused by prenatal cannabinoid exposure. It is important to note that many of the studies that assess offspring growth describe a reduction in body weight at birth followed by rapid catch-up growth^11, 18, 45–47^. Growth restriction followed by postnatal catch-up growth are associated with long-term consequences related to metabolic and cardiac health in adolescent and adult life^48, 49^. Currently, there is a decreased perception of risk regarding regular cannabis use which, in part, contributes to the rising rates of cannabis consumption across all population demographics, including pregnant women^50^. However, our study along with other preclinical studies provide convincing evidence that prenatal cannabinoid exposure does alter fetal and placental developmental programming that has the potential to translate to offspring development.

## Author contributions

R.C.W. and M.N.R. contributed to the experimental design. A.H.C. performed dosing of all animals. A.H.C., C.R.C., P.D., C.P. and R.C.W. performed tissue collections, sample processing, and data analysis. R.C.W. wrote the first draft of the manuscript and all authors assisted with editing of the manuscript.

